# Mapping bias overestimates reference allele frequencies at the *HLA* genes in the 1000 Genomes Project phase I data

**DOI:** 10.1101/013151

**Authors:** Débora Y. C. Brandt, Vitor R. C. Aguiar, Bárbara D. Bitarello, Kelly Nunes, Jérôme Goudet, Diogo Meyer

**Affiliations:** Department of Genetics and Evolutionary Biology, University of São Paulo, 05508-090 São Paulo, SP, Brazil; Department of Ecology and Evolution, Biophore, University of Lausanne, CH-1015 Lausanne, Switzerland

**Author notes:** **Corresponding author** Diogo Meyer, Departamento de Genética e Biologia Evolutiva, Rua do Matão, 277, São Paulo, SP 05508-090, Brazil.

**Keywords:** NGS, mapping bias, 1000 Genomes, HLA

## Abstract

Next Generation Sequencing (NGS) technologies have become the standard for data generation in studies of population genomics, as the 1000 Genomes Project (1000G). However, these techniques are known to be problematic when applied to highly polymorphic genomic regions, such as the Human Leukocyte Antigen (*HLA*) genes. Because accurate genotype calls and allele frequency estimations are crucial to population ge-nomics analises, it is important to assess the reliability of NGS data. Here, we evaluate the reliability of genotype calls and allele frequency estimates of the SNPs reported by 1000G (phase I) at five *HLA* genes (*HLA-A*, *-B*, *-C*, *-DRB1*, *-DQB1*). We take advantage of the availability of *HLA* Sanger sequencing of 930 of the 1,092 1000G samples, and use this as a gold standard to benchmark the 1000G data. We document that 18.6% of SNP genotype calls in *HLA* genes are incorrect, and that allele frequencies are estimated with an error higher than ±0.1 at approximately 25% of the SNPs in *HLA* genes. We found a bias towards overestimation of reference allele frequency for the 1000G data, indicating mapping bias is an important cause of error in frequency estimation in this dataset. We provide a list of sites that have poor allele frequency estimates, and discuss the outcomes of including those sites in different kinds of analyses. Since the *HLA* region is the most polymorphic in the human genome, our results provide insights into the challenges of using of NGS data at other genomic regions of high diversity.

**Data available in public repositories**

https://github.com/deboraycb/reliability_hla_1000g

## INTRODUCTION

Whole genome resequencing data for large numbers of human individuals, as generated by the 1000 Genomes Project (www.1000genomes.org), provide unprecedented amounts of information about microevolutionary processes and demographic histories. Such inferences rely on either genotypic or allelic frequency information for each variable position, which constitute the data for downstream analyses and hypothesis testing.

The calling of SNPs and genotypes and the estimation of allele frequencies from Next Generation Sequencing (NGS) has undergone rapid development, along with likelihood-based and Bayesian methods created to deal with challenges associated to heterogeneity in read quality and coverage (Nielsen et al. 2011). In Phase I of the 1000 Genomes Project, genotypes were called using a combination of different approaches: first, primary call sets were independently generated by different centers with different sequencing platforms, alignment and variant calling methods; then, a consensus SNP call set was generated and made publicly available (The 1000 Genomes Project Consortium 2012).

The data generated by the 1000 Genomes Project have frequently been used to make inferences about evolutionary processes affecting our species, including the detection of targets of natural selection (Hernandez et al. 2011; Ward and Kellis 2012; Andersen et al. 2012) and understanding the genetic basis of complex phenotypes (Lappalainen et al. 2013). In addition, the detailed catalogue of genetic variation it provides across multiple human populations has been used to understand the processes affecting specific genes.

Among the well documented targets of selection is the Major Histocompatibility Complex (MHC) region of the human genome, which harbors the highly polymorphic classical Human Leukocyte Antigen *(HLA)* class I and II loci. The interest in these loci stems from their strong association to various autoimmune disorders (Sollid et al. 2014), susceptibility and resistance to infection (Chapman and Hill 2012), and striking signatures of genetic variation indicating strong balancing selection (Meyer and Thomson 2001). Such types of investigations can naturally be extended to the analysis of the 1000 Genomes data, which provide a rich resource of population genetic variation within and around *HLA* genes.

In spite of this interest, the use of NGS data for *HLA* loci is hampered by a major technical hurdle, which is the mapping of short sequence reads to genes that are both highly polymorphic and which constitute a multi-gene family. The high polymorphism may decrease the probability that short reads will be successfully mapped to the reference genome, in the event that the sequenced individual carries a variant that is highly diverged from that used in the index (Nielsen et al. 2011). In addition, the fact that many *HLA* genes have close paralogues increases the chance that a read will map to two or more genomic regions, leading it to be discarded from most sequencing analyses pipelines, and thus decreasing the amount of usable information for genotype calling (Treangen and Salzberg 2012).

Previous studies explored the applicability of NGS to genotype the *HLA* alleles of an individual, where an allele is typically defined as the haplotype determined by a combination of SNPs within a given *HLA* gene (e.g. (Erlich et al. 2011; Major et al. 2013)). To this end, Erlich et al. (2011) proposed NGS methodologies in which different steps - from sample preparation to haplotype level allele calling - were adapted to deal with the issues of high polymorphism and paralogy of *HLA* genes. In this way, they were able to successfully validate their methodology in a study of 270 samples that had been previously typed by sequence specific oligonucleotide (SSO) hybridization, which they treated as a gold standard dataset. The same gold standard dataset was used by Major et al. (2013), who also examined the reliability of calling *HLA* alleles using NGS, but using the 1000 Genomes alignment data, and showed that this publicly available dataset can be used for this purpose, after appropriate filters (e.g. coverage) are applied.

Both Erlich et al. (2011) and Major et al. (2013) were interested in using NGS data to determine *HLA* alleles. Information regarding *HLA* alleles is of biomedical relevance since *HLA* genotypes are often an important covariate to account for in association studies, and *HLA* typing is critical to hematopoietic transplantation. In this study, however, we evaluate the quality of SNP level genotype calls from the 1000 Genomes at the *HLA* genes.

The analysis of genotype and allele frequencies for SNPs contained within *HLA* genes has proven of great value in biomedical and evolutionary studies, and the 1000 Genomes dataset is a recurrently used resource in this context. Examples of the use of *HLA* SNP data from the 1000 Genomes Project include: (a) In GWAS studies SNPs in *HLA* genes are often associated with phenotypes of interest, and it is useful to understand the prevalence of these variants in additional populations; (b) GWAS studies benefit from knowledge of the haplotype structure surrounding *HLA* genes, which can be inferred from the dense SNP data of the 1000 Genomes for multiple populations (e.g. Hill-Burns et al. 2011); (c) When testing for selection, many studies have found strong evidence associated to the *HLA* region, using the 1000 Genomes as a source of polymorphism data (e.g. Leffler et al. 2013).

All the above applications of the 1000 Genomes Project SNP data in *HLA* genes are dependent on the reliability of genotype calls at each SNP. However, no study to date has provided a detailed survey of the reliability of individual genotype calls and allele frequency estimates at the SNPs in *HLA* genes, in spite of their frequent usage. We address this issue, discuss likely causes for cases of incorrect genotype calls and provide a list of reliable sites for the *HLA* loci in the 1000 Genomes data. As in previous studies (Erlich et al. 2011; Major et al. 2013), we used a dataset in which individuals had their *HLA* genes genotyped using Sanger sequencing as a gold standard to benchmark the genotypes called at the 1000 Genomes Project. However, differently from these other studies, which were interested in reconstructing the *HLA* haplotypes using NGS, here we have deconstructed the haplotypes determined from Sanger sequencing data into SNPs, and compared genotypes at the SNP level to the 1000 Genomes data. We took advantage of the recent availability of a dataset of Sanger sequencing based *HLA* genotyping of *HLA-A*, *-B*, *-C*, *-DQB1* and *-DRB1* for 930 of the samples from the 1000 Genomes Project (Gourraud et al. 2014). Our results have implications for other studies that use SNP data from the 1000 Genomes in order to estimate allele frequencies. Because *HLA* loci are the most polymorphic in the human genome, they most likely represent the worst case scenario for mapping bias and, consequently, allele frequency estimation error.

## METHODS

In this study we compare NGS genotype calls and allele frequency estimates reported by the 1000 Genomes Project with those obtained in a study which used Sanger sequencing to genotype *HLA* genes. For the purpose of our analysis we assembled a dataset comprising the intersection of the 1000 Genomes and Sanger sequencing samples, resulting in 930 individuals from 12 populations. Figure 1 summarizes the pre-processing of both datasets, which preceded genotype and allele frequency comparisons.

### 1000 Genomes dataset (1000G)

SNP genotypes were acquired from the chromosome 6 integrated Variant Call Format (VCF) file from version 3 of the 1000 Genomes Project Phase I data, which is available at ftp://ftp.1000genomes.ebi.ac.uk/vol1/ftp/release/20110521/ (The 1000 Genomes Project Consortium 2012). We selected only SNPs in exons encoding the antigen recognition sites (ARS), which are exons 2 and 3 for *HLA-A,-B*, and *-C* (Bjorkman et al. 1987) and exon 2 for *HLA-DQB1* and *-DRB1* (Brown et al. 1993). Sites were selected based on the most inclusive coordinates of the RefSeq database in 22 July 2014 (see File S1). Both SNP and sample selection were carried out using VCFtools v0.1.12a (Danecek et al. 2011).

### HLA reference panel by Gourraud et al. (2014) (PAG2014)

Gourraud et al. (2014) typed class I *HLA-A*, *-B* and *-C*, and class II *HLA-DRB1* and *-DQB1* genes of 1266 individuals from 14 different populations in Africa, Europe, Asia and America. The *HLA* sequence-based typing was performed with specific PCR amplification of ARS exons followed by Sanger sequencing. Data are available at the dbMHC website (http://www.ncbi.nlm.nih.gov/gv/mhc/xslcgi.fcgi?cmd=cellsearch) (Helmberg et al.).

Data from Gourraud et al. (2014) are available in the form of *HLA* allele names per individual. Allele naming for *HLA* genes follows specific rules (Marsh et al. 2010). Briefly, allele names are composed of a letter indicating the locus, followed by 2 to 4 numeric fields separated by colons. Each numeric field indicates specific forms of variation: the 1st field distinguishes groups of alleles by serological type, and the following fields distinguish nonsynonymous polymorphisms, synonymous polymorphisms, and non-coding differences, respectively. In order to obtain SNP genotypes and frequencies from the Sanger sequencing data, we converted all allele names to their associated sequences for ARS encoding exons. Sequences were acquired from the IMGT database (Robinson et al. 2013), which keeps a well curated repository of all known *HLA* allele sequences.

Our analysis was restricted to ARS exons because the *HLA* typing method used by Gourraud et al. (2014) only probed genetic variation in these specific exons. As a consequence, multiple *HLA* alleles are compatible with the sequencing results, since the sites that differentiate them are in other exons. This results in what we refer to as an “ambiguous allele call” for an *HLA* allele (e.g., the allele is identified as B*35:03 but we cannot establish whether it is B*35:03:01 or B*35:03:02, or a group of alleles is attributed to an individual, such as B*35:02/B*35:03/B*35:04). Ambiguous allele calls may also happen when sequencing has low quality at bases that differentiate two alleles. In addition, there are also genotypic ambiguities, which occur when different pairs of alleles are compatible with the sequencing results. For individuals that bear ambiguous alleles, we created a consensus sequence in which ambiguous sites were reported with both possible alleles (e.g. A/T, see Figure 1). In this way, we incorporate the uncertainty associated to the sequence-based typing into downstream analyses.

Although we cannot rule out technical errors in the Sanger sequencing that generated the PAG2014 data (Gourraud et al. 2014), we assume that this method provides the most reliable estimate of *HLA* alleles (and hence SNP genotypes), and will serve as a standard to estimate the reliability of genotype calls and allele frequencies for the 1000 Genomes data (De Santis et al. 2013).

**Figure 1:**
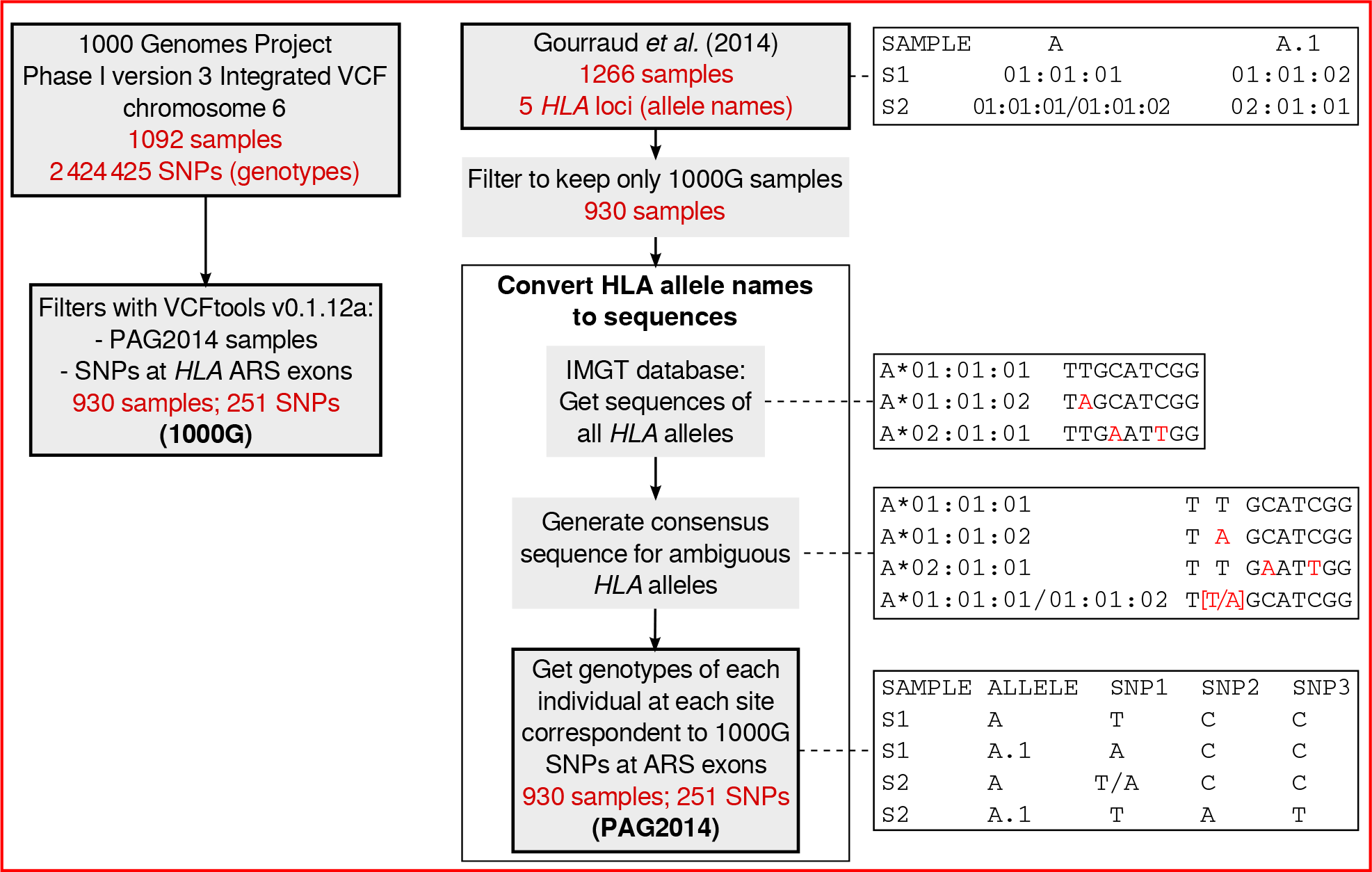
Workflow for preparation of next generation sequencing dataset from the 1000 Genomes Project (1000G) and Sanger sequencing dataset generated by Gourraud et al. (2014) (PAG2014) for comparisons of genotypes and allele frequencies (see main text).

### Genotype comparisons

We initially quantified how well the 1000G and PAG2014 data agreed with respect to genotype calls. Genotypes at each site in each individual were compared between the 1000G data and the PAG2014 data, here considered as a gold standard. In the case of sites with ambiguity (e.g. T/A) in the PAG2014 data, if one of the two possible alleles matched an allele present in the 1000G, we considered this an allele match and PAG2014 was corrected, by attributing the allele present in the 1000G data to the ambiguous site. After correcting the ambiguous sites in PAG2014, we only considered genotypes to be a match if both alleles in 1000G were present in the PAG2014 data, at that site.

Throughout this paper, sites are numbered according to their position in the ARS exons coding sequences (1-546 at the class I loci and 1-270 at the class II loci).

### Allele frequency comparisons

After correcting all possible ambiguities in PAG2014 (as described above), we calculated allele frequencies for SNPs in both datasets. By comparing the frequency of the reference allele in 1000G to its value in PAG2014 we assessed the accuracy of allele frequency estimation. The reference allele was defined as the allele present in the hg19 build of the reference sequence of the human genome. RefSeq IDs of the reference sequences used for each *HLA* gene are reported on File S1.

We computed the error in 1000G frequency estimates per site *i (FE_i_)* as:

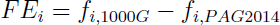

where *f_i;1000G_* and *f*_*i;PAG*2014_ are the frequency of the reference allele at site *i* in 1000G and PAG2014, respectively. We also computed the mean absolute error in frequency estimates per gene as a mean of absolute *FE_i_* for all sites within a gene (MAE):

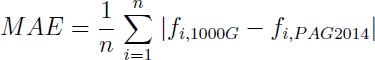

where *n* is the number of SNPs in the gene.

### Coverage in 1000G

Sequencing coverage per individual per site was calculated from the 1000 Genomes Project phase I BAM files for the low coverage experiments. Using the.genomeCoverageBed program from BEDTools (Quinlan and Hall 2010). BAM files are available on ftp://ftp.1000genomes.ebi.ac.uk/vol1/ftp/phase1/data/[sampleID]/alignment/. Only low coverage BAM files were used to estimate coverage because genotype likelihoods for the data we analyzed (1000 Genomes Project Phase I integrated VCF files) were estimated from this source. Genotype likelihoods were estimated from high coverage exome BAM files only for a minority of sites that were exclusively discovered on the exome experiments, and were not used in the coverage analysis (See Table S1).

### Testing for mapping bias

After demonstrating that there is an overestimation of reference allele frequency in the 1000G SNPs (see Results), we hypothesized that mapping bias was the underlying cause. To test this hypothesis we examined whether reads carrying the alternative allele at a SNP are less likely to map to the reference genome than reads carrying the reference allele. First, for each *HLA* allele present in the PAG2014 dataset, we defined windows of 51 basepairs that were centered on each SNP (25 basepairs upstream and 25 basepairs downstream of the SNP, including non-polymorphic sites). The set of windows centered on a specific SNP was then separated in two groups: i) those that carry the reference allele at the central site and ii) those that carry the alternative allele at the central site. Next, all windows were compared to the reference genome (hg19) sequence, and the number of mismatches was counted, excluding the mismatch at the central SNP. If mapping bias was influencing allele frequency estimates, we expected that, for SNP positions with overestimation of the reference allele frequency in the 1000G, the alternative alleles would be flanked by additional alternative alleles (and thus have a higher mismatch count against the reference sequence).

## RESULTS

### Genotypic mismatch frequency

We found that, on average, 18.6% of genotypes where mismatched between 1000G and PAG2014 when individual genotypes for each site in the 5 classical *HLA* genes were compared. *HLA-B*, *-DQB1* and *-DRB1*, which are the *HLA* genes with the highest levels of nucleotide diversity (Buhler and Sanchez-Mazas 2011), also show the highest proportion of genotype mismatches (23%, 21% and 27%, respectively). We also observed that mismatches are specially concentrated on a few sites (Figure 2), with 18.7% of sites concentrating 50% of the mismatches over the 5 loci we analyzed.

**Figure 2:**
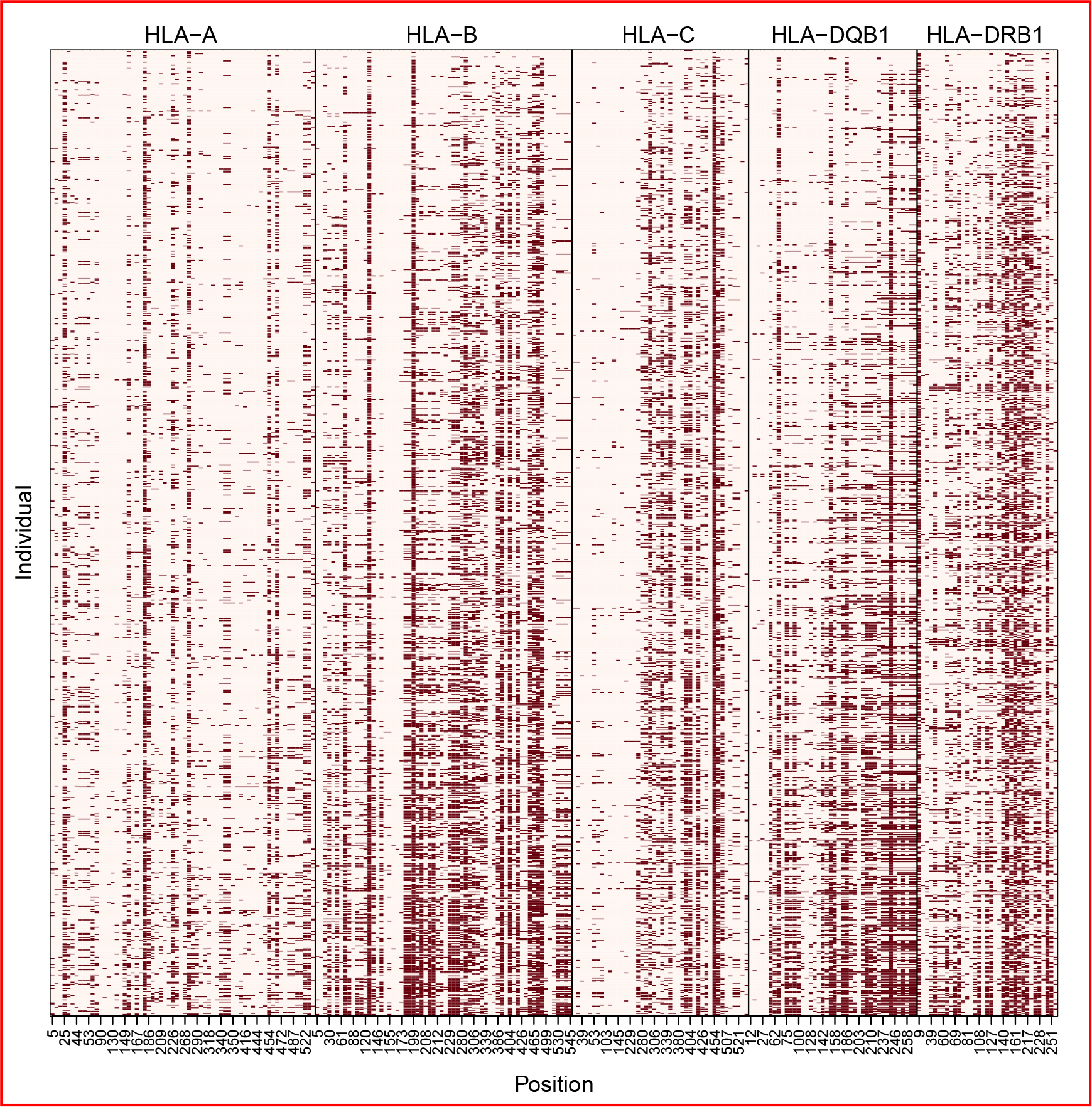
Genotype mismatches between the 1000G and PAG2014 datasets. Results per polymorphic site (“Position”) and per individual (930 in total). Individuals are ordered by number of mismatches (individuals with less mismatches on top). Sites are numbered according to their position in ARS exons coding sequence. Dark squares indicate mismatches between genotypes in the two datasets.

### Reference allele frequency accuracy

Accuracy of estimation of allele frequencies in 1000G was assessed comparing the observed frequency of the reference allele in the 1000G data with that of PAG2014, for both the global dataset (consisting of a pooled set of all individuals) and for each population separately (see Figure S1-S5). We chose a difference of 0.1 between the frequencies on both datasets as a threshold that determines a “large frequency difference”.

For the global dataset (Figure 3) we found that for *HLA-A* and *-C* most SNPs have similar frequency estimates for 1000G and PAG2014, with few large deviations (only 9/66 and 8/44 SNPs with absolute difference in frequencies (|*FE*|) larger than 0.1, respectively). The *HLA-DQB1* locus shows an intermediate proportion of SNPs with large deviations (10/42 SNPs with |*FE*| > 0.1), and *HLA-B* and *HLA-DRB1* show the greatest proportion of sites with large frequency differences between 1000G and PAG2014 (23/64 and 15/35 sites with |*FE*| > 0.1). Overall, the mean absolute difference in frequency between SNPs in the 1000G and PAG2014 data is 0.08, and it is higher at the *HLA* genes with the highest levels of nucleotide diversity (*HLA-B*, *-DQB1* and *-DRB1* all deviate by ±0.1).

**Figure 3:**
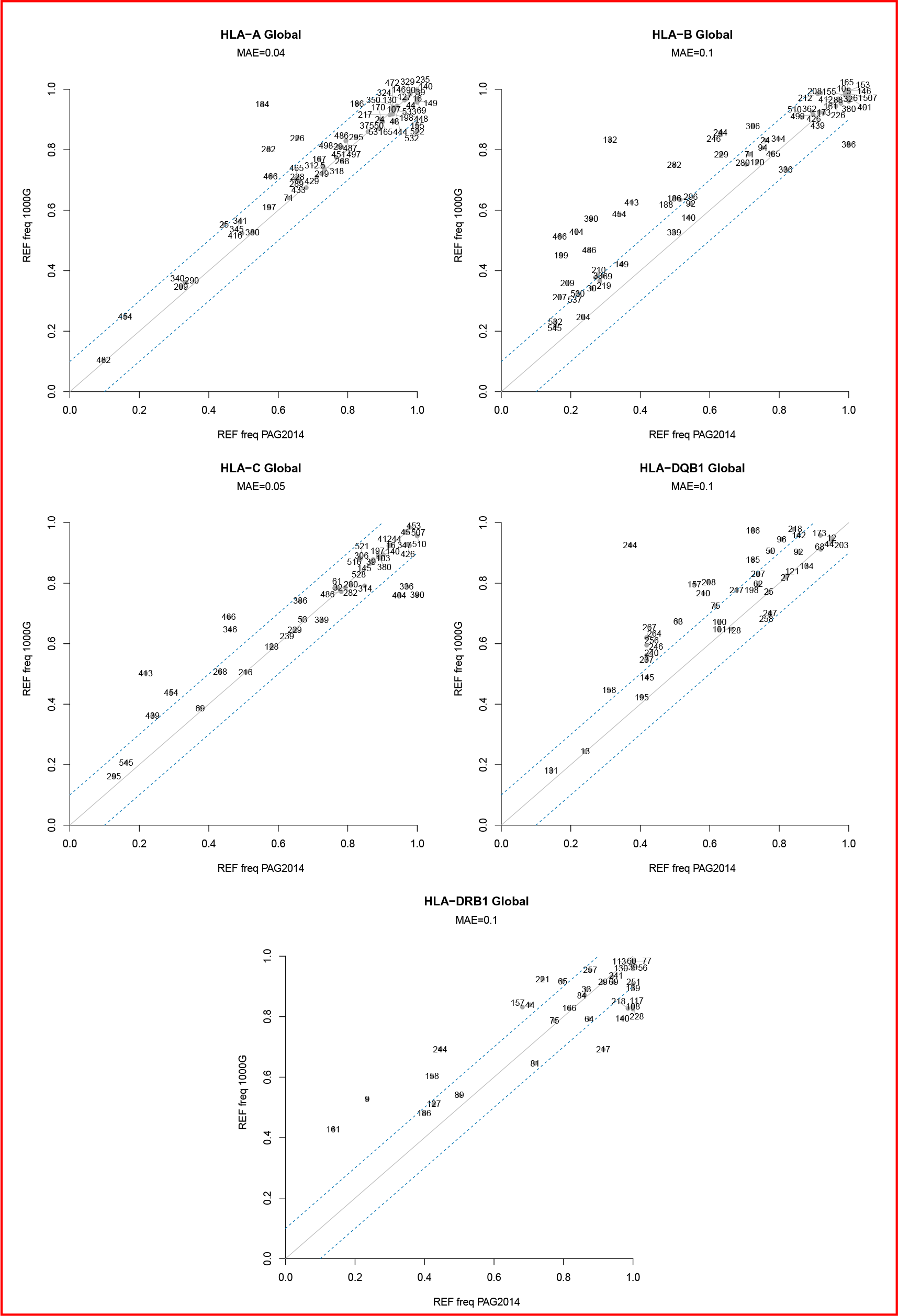
Reference allele frequency per site in each *HLA* gene in the 1000 Genomes (1000G) and Sanger sequencing (PAG2014) datasets. Continuous line indicates the expected relationship (i.e., no difference) between 1000G and PAG2014. Dashed lines indicate a ±0.1 deviation from the expected frequency (as estimated from PAG2014 dataset). MAE (mean absolute error) defined in Methods. Numbers indicate site position in ARS exons sequence.

The proportion of genotype mismatches and allele frequency deviations per site are highly correlated (Pearson correlation = 0.86, p-value < 10^-16^; Figure S6). However, some SNPs with a high proportion of genotype mismatches have well estimated allele frequencies. One example is site 465 at *HLA-B*, in which 44% of genotypes are mismatched, but |*FE*| is only 0.007. Overall, 15 sites have more than 25% mismatched genotypes while showing |*FE*| < 0.1. (See Figure S6). This is possible when the frequency of genotype errors in which the reference allele is overrepresented is similar to the frequency of errors in which the alternative allele is overrepresented.

Allele frequency at the Affymetrix Axiom array: Because genotyping arrays constitue an additional frequently used resource to genotype SNPs within *HLA* genes, playing an important role in GWAS studies, we have also investigated the accuracy of allele frequency estimation from this genotyping technology. We estimated allele frequencies from Affymetrix Axiom array data, and we found that those allele frequency estimates are as reliable as the ones from the 1000 Genomes NGS data, at the same SNPs (see Figure S7).

### Relationship between sequencing coverage and genotypic mismatches

To investigate whether low sequencing coverage could explain genotype mismatches and deviations from expected allele frequencies, we compared sequencing coverage between mismatched and matched genotypes (Figure 4a) and assessed the relationship between coverage and frequency deviation (Figure 4b).

**Figure 4:**
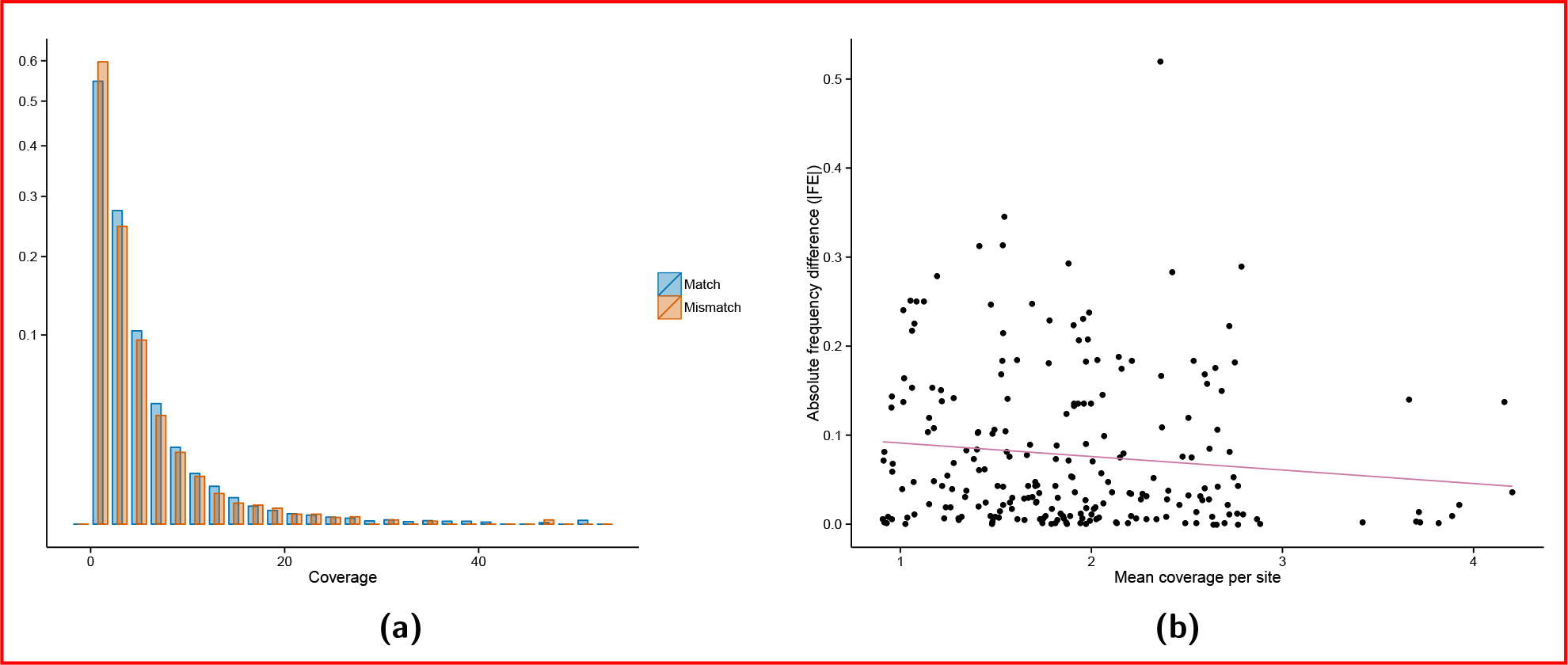
(a) Distribution of coverage (x-axis) at matched and mismatched genotypes; y-axis is the square root of the relative frequency (Mann-Whitney one-tailed test p-value < 10^−16^); (b) Relationship between mean coverage (x-axis) and absolute frequency difference (|*FE*|, y-axis) between 1000G and PAG2014 (r = -0.11, p-value = 0.09). All polymorphic sites from *HLA-A*, *-B*, *-C*, *-DRB1* and *-DQB1* genes are included in both a and b.

Sites with mismatched genotypes have on average lower sequencing coverage than sites with matched genotypes (Figure 4a; Mann-Whitney one-tailed test p-value < 10^-16^). This is the expected relationship if low sequencing coverage explains genotype mismatches between datasets. However, the difference in sequencing coverage between sites with matched and mismatched genotypes is small (mean coverage in matching genotypes is 1.95, and 1.75 in non-matching genotypes, a difference of 6,2%), and has likely achieved very high significance only due to the large number of observations. Similarly, correlation between allele frequency deviation and sequencing coverage is weak and not significant (Figure 4b; r = -0.11, p-value = 0.09), although the direction of correlation is in agreement with what would be expected if lower coverage explained larger deviations in frequency estimation. We therefore investigated other factors that may account for errors in genotype calling.

### Direction of frequency deviation

Most of the deviations in allele frequency estimates are in the direction of an overestimation of reference allele frequencies in the 1000 Genomes data (Figure 3). This information is summarized in figure 5 which shows the location and magnitude of deviations between the 1000 Genomes and PAG2014 data.

**Figure 5:**
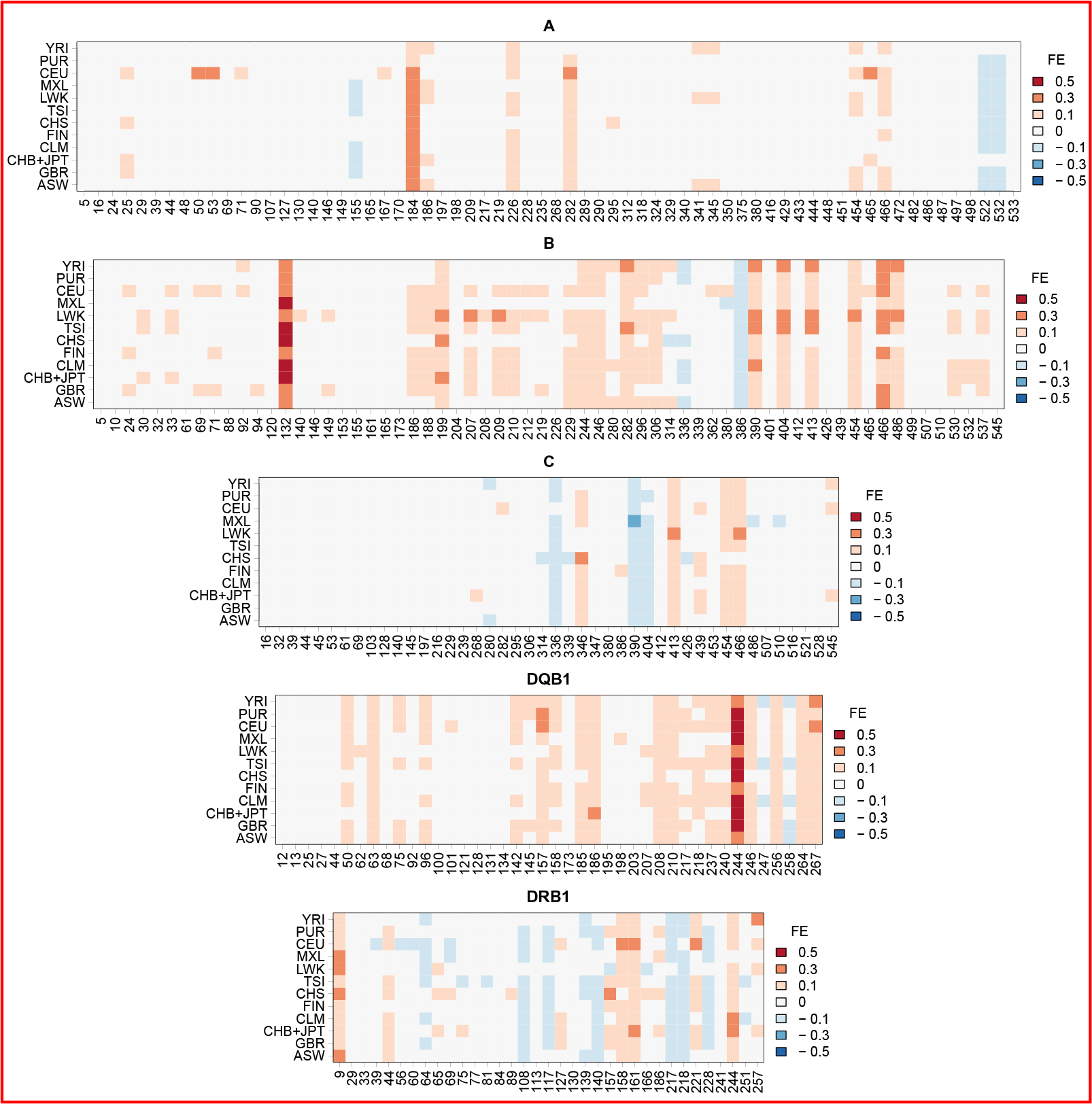
Difference in reference allele frequency between 1000G and PAG2014, measured by *FE* (see Methods), at each polymorphic site, in each population. Shades of red indicate overestimation of reference allele frequency and shades of blue indicate underestimation of reference allele frequency in 1000G. Full population names are given in Table S2.

The overall shift in the direction of overestimating reference alleles is summarized in Table 1, which shows the number of SNPs with more than 0.1 frequency difference in at least two populations, for each locus. For *HLA-A*, *-B* and *-DQB1* most sites with large frequency differences between 1000G and PAG2014 are skewed in the direction of overestimating the reference allele (p-value = 0.057 for *HLA-A* and p-value < 10^-4^ for *HLA-B* and *DQB1*, binomial test for null hypothesis of equal numbers of deviations in direction of REF or ALT), whereas *HLA-C* and *HLA-DRB1* show no evidence for an excess of large deviations in the direction of reference alleles.

**Table 1:**
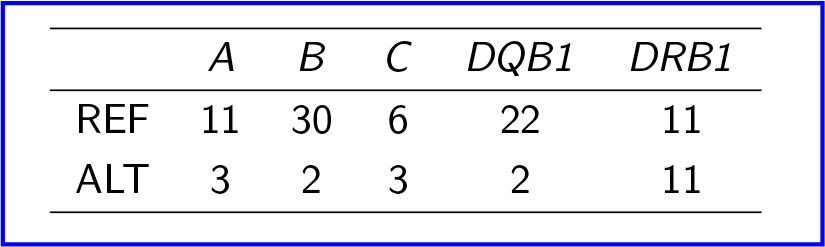
Number of sites with overestimation of reference (REF) or alternative (ALT) allele frequency in each *HLA* locus (|*FE*| > 0.1 in 2 or more populations). Genomic coordinates of those sites are given in Table S3.

|  | *A* | *B* | *C* | *DQB1* | *DRB1* |
| --- | --- | --- | --- | --- | --- |
| REF | 11 | 30 | 6 | 22 | 11 |
| ALT | 3 | 2 | 3 | 2 | 11 |

### Testing for mapping bias

We hypothesized that the observed reference allele bias was caused by a lower efficiency in the mapping of reads containing the alternative allele. This is expected under the assumption that the reads carrying the alternative allele on average have more differences with respect to the reference genome (used by the 1000 Genomes Consortium as the index to align NGS reads) than reads carrying the reference allele. In this scenario, some sites would have a stronger bias than others if the alternative alleles in those sites are flanked by additional alternative alleles.

To test this hypothesis, we aligned sequences of all alleles present in PAG2014 to the reference genome, and defined windows of 51 base pairs around each SNP. We then quantified the number of differences with respect to the reference genome for windows surrounding i) reference (REF) and ii) alternative (ALT) alleles. If reference allele mapping bias is driving errors in frequency estimation, it is expected that sites with an overestimation of reference allele frequency would present the following pattern: windows carrying the reference allele (REF) with fewer differences to the reference genome than sequences centered on the alternative allele (ALT). For sites with well estimated frequencies, on the other hand, we did not expect such a difference between REF and ALT windows.

To illustrate this effect, Figure 6 shows the results for the two most extreme cases of frequency deviation shown in Figure 5: site 244 of *HLA*-DQB1 and site 132 of *HLA*-B (0.56 and 0.52 absolute increase in reference allele frequency in the 1000 Genomes data with respect to PAG2014). In both cases, ALT windows bear more differences to the reference sequence than REF windows.

**Figure 6:**
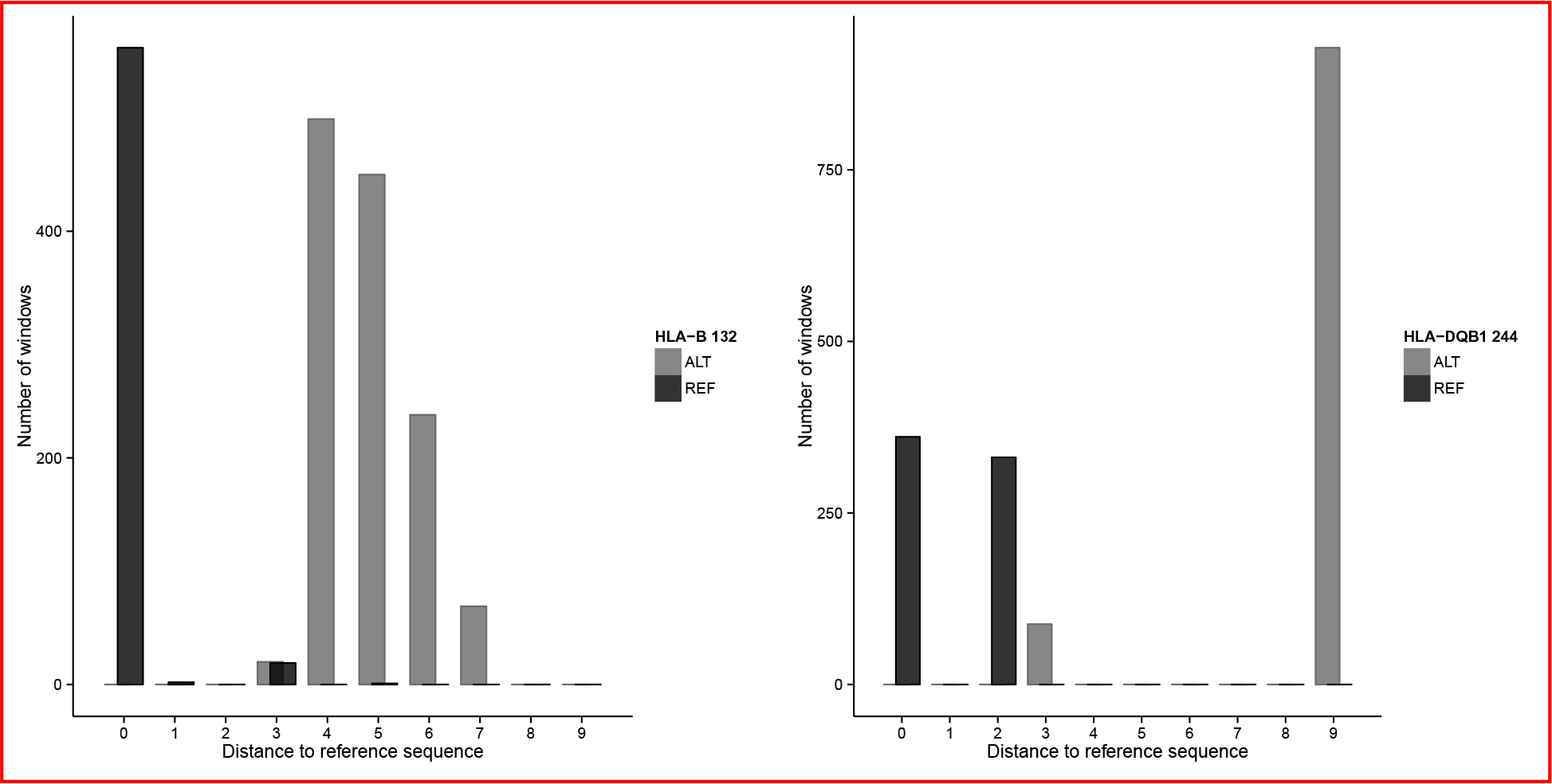
Number of differences to the reference genome at 1860 51bp windows centered at sites *HLA-B* 132 and *HLA-DQB1* 244 with reference (REF) or alternative (ALT) allele at those sites. Windows were defined from all *HLA* alleles present in the 930 samples from the PAG2014 dataset.

These results support the hypothesis that these sites with poorly estimated allele frequencies have their alternative alleles residing in haplotypes with substantially more differences with respect to the reference genome than haplotypes centered on the reference allele, thus accounting for the observed bias.

To gain a broader perspective of this issue we classified SNPs from the *HLA* loci with reference allele bias (*HLA-A*, *-B*, and *-DQB1*) into three categories: i) sites where the reference allele frequency was overestimated, i.e. *FE* > 0.1 (“overestimated”), ii) sites where the reference allele frequency was underestimated, i.e. *FE* < -0.1 (“underestimated”) and iii) sites where allele frequencies were well estimated (|*FE*| < 0.01, here refered to as “well estimated”). We compared these three categories of sites with respect to the number of differences relative to the reference genome in REF and ALT windows (Figure 7). We found that the overestimated group has significant excess of differences at alternative allele bearing haplotypes. In this group of SNPs, ALT windows have on average 4.4 other differences relative to the reference genome, while those centered on the reference allele (REF) have 1.9 differences (excess of differences on windows centered on the alternative allele was tested with a one tailed Mann-Whitney test; p-value < 10^-16^). Sites with well estimated or underestimated reference allele frequency, on the other hand, do not show a similar excess of differences in the haplotypes bearing the alternative allele, although the difference between REF and ALT windows is statistically significant due to the large sample size (well estimated: ALT mean=1.7; REF mean=1.8; one tailed Mann-Whitney test p-value < 10^-16^; underestimated: ALT mean = 1.9; REF mean = 1.2; one tailed Mann-Whitney p-value < 10^-16^).

**Figure 7:**
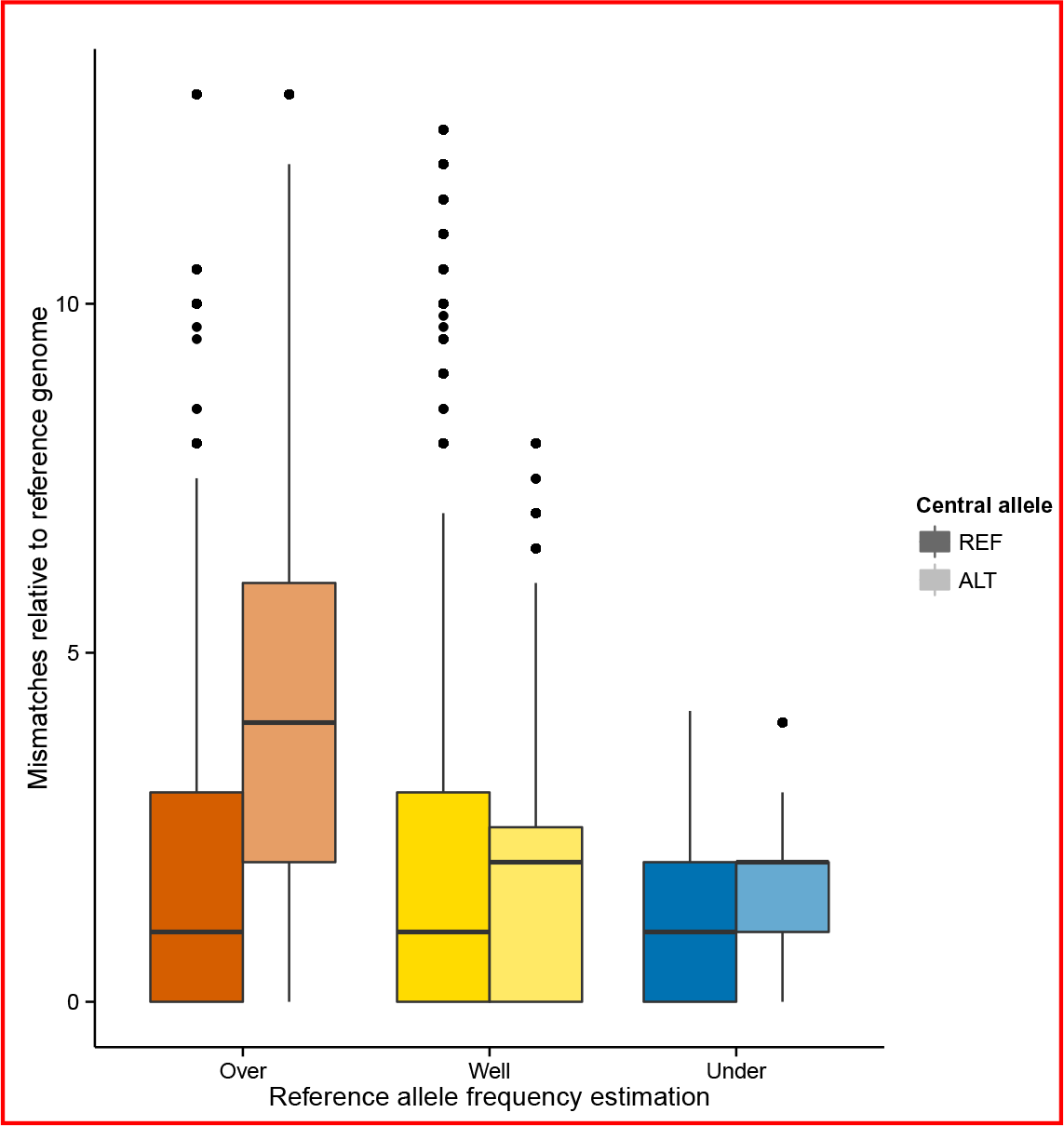
Number of differences to the reference genome at 51bp windows centered at each SNP in the *HLA-A*, *-B* and *-DQB1* genes. Windows around each SNP were defined from the set of 1860 alleles present in the 930 samples from the PAG2014 dataset. Next, the set of windows was divided in three groups: those centered on SNPs with overestimated, well estimated and underestimated reference allele frequencies (red, yellow and blue boxplots, respectively). Then, each group was divided in two: windows in which the central site contains the reference allele (*REF*, dark boxplots) and windows centered on an alternative allele (*ALT*, light colored boxplots). Upper and lower hinges correspond to the 25th and 75th percentiles, horizontal lines represent the median, whiskers are 1.5 times the interquartile range, and outliers are represented by dots.

### Impact of biases in frequency estimation to population genetic statistics

Our analysis was able to identify a subset of SNPs in the *HLA* genes for which genotype calls and allele frequency estimates from the 1000G showed a high error rate with respect to the PAG2014 dataset. To evaluate the impact of the errors introduced by including these sites in population genetic analyses, we compared the distribution of sample heterozygosity between the sites with low and high error rates. Heterozygosity is defined as *H* = 2*p*(1 - *p*) for biallelic loci, as is the case for the 1000 Genomes Phase I SNPs, since tri-or quad-allelic SNPs were not reported on Phase I.

The removal of sites with poor frequency estimates (|*FE*| > 0.1 in at least two populations) results in a marked change in the distribution of *H*, with a significant drop in the frequency of sites with large *H* and a shift in the distribution towards lower values (Figure 8). Note that the values in the Figure 8a are for *H* values estimated from the PAG2014 data, implying that the high values of *H* among “excluded” sites are not due to the deviations in allele frequencies generated by NGS errors, but is the true heterozygosity at those sites. These results therefore document that because sites with high heterozygosity tend to have greater deviations from the “true” frequency (i.e., based on the PAG2014 dataset), the removal of poorly estimated sites results in a reduction in *H* values.

**Figure 8:**
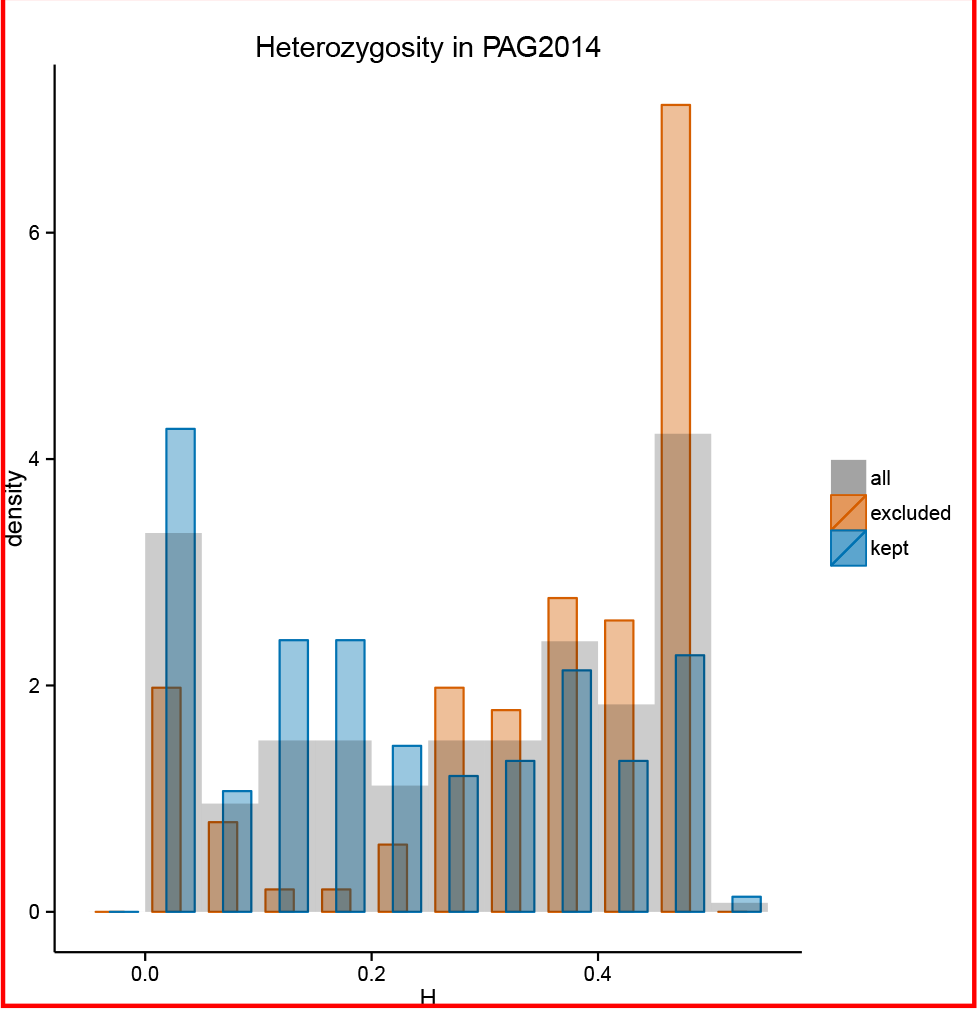
Heterozygosity of SNPs at *HLA* genes estimated from the PAG2014 dataset. Orange bars show distribution of heterozygosity at sites with a high error rate in frequency estimation (|*FE*| > 0.1 in two or more populations). Blue bars show the distribution of heterozygosity after exclusion of SNPs with high error rate.

### The effect of heterozygosity on allele frequency estimation bias

We found an overall positive correlation between SNP heterozygosity and the magnitude of error in allele frequency estimates (Figure 9a; Pearson’s correlation = 0.32; p-value = 1.938 × 10^-7^). This result provides further evidence that sites with higher heterozygosity tend to have poorer estimates for allele frequencies in the 1000G. Also, heterozygosity is even more strongly correlated to the deviation in frequency, considering the direction of the deviation (Figure 9b; Pearson’s correlation = 0.59; p-value < 10^-16^). Together, these results show that *HLA* SNPs with higher heterozygosities not only have more errors in frequency estimation but also a stronger bias towards overestimation of reference allele frequency.

**Figure 9:**
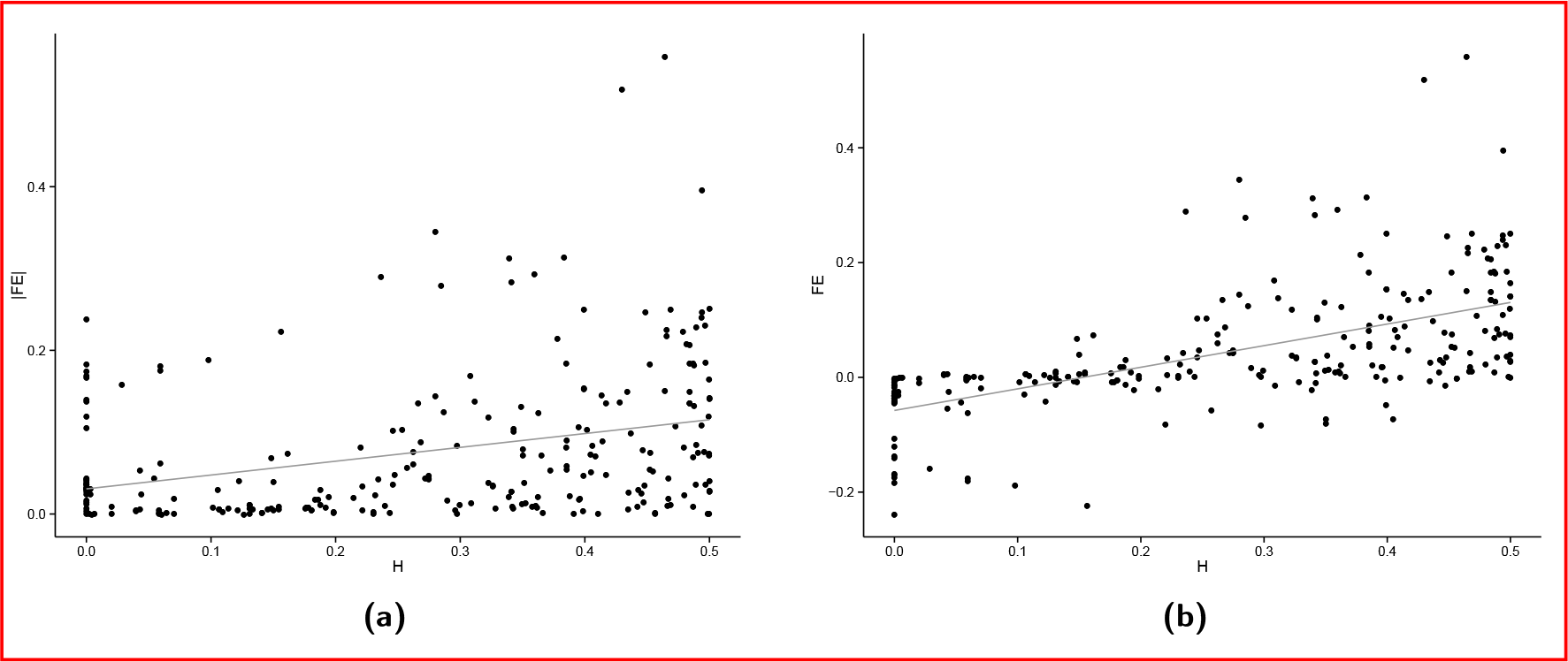
Relationship between SNP heterozygosity (H) and (a) absolute value of deviation (|*FE*|; Pearson’s correlation = 0.32; p-value = 1.938 × 10^-7^) or (b) magnitude and direction of deviation (*FE*; Pearson’s correlation = 0.59; p-value < 10^-16^).

## DISCUSSION

The 1000 Genomes Project data were generated by various sequencing centers, which relied on different sequencing platforms, read lengths, aligners and variant and genotype calling algorithms (The 1000 Genomes Project Consortium 2012), creating challenges to an overall assessment of data reliability. In this study, we specifically examine the performance of NGS based genotype calls and allele frequency estimates for the highly polymorphic and intensely studied classical *HLA* genes. We took advantage of the possibility of comparing downstream genotype calls from the 1000 Genomes and *HLA* typing based on Sanger sequencing for the same set of samples to assess data quality and test hypothesis about possible biases.

We show that the 1000 Genomes SNPs called in the *HLA* genes have many differences at the genotype level, when compared to results obtained using Sanger sequencing. However, considerably high genotype mismatching is possible with only modest deviations in allele frequencies, and we conclude that for the 1000 Genomes data allele frequency estimates for SNPs at *HLA* genes are considerably more reliable than the individual genotype calls.

The errors in frequency estimates in the 1000 Genomes NGS data are biased towards an overestimation of the reference allele frequency, a pattern consistent with read mapping bias. Mapping bias is well known for NGS, and highly polymorphic regions such as *HLA* genes are particularly susceptible to its effects (Nielsen et al. 2011). In our study, *HLA-A*, *-B*, and *DQB1* show evidence of reference allele mapping bias. The *HLA-DRB1* locus, on the other hand, did not present reference allele frequency overestimation, a finding that can be explained by the existence of multiple copies of this gene (both pseudogenes and functional copies), which may result in biases/errors that make reference allele bias comparatively less visible (Degner et al. 2009). The *HLA-C* locus also shows a weaker reference allele bias, a pattern that may be explained by its lower degree of polymorphism which leads to a decrease in the number of mismatches of reads with respect to the reference genome, thus decreasing the mapping bias.

We provide a list of unreliable SNPs within the *HLA* genes, defined by us as those with an absolute difference in frequency larger than 0.1 (|*FE*| > 0.1) in two or more populations (Table S3). We show that these unreliable SNPs on average have higher heterozygosities in our gold standard dataset. As a consequence, although filtering out those unreliable sites improves the overall accuracy in allele frequency estimation, it leads to an underestimation of the mean heterozygosity of SNPs in *HLA* genes, a bias that should be taken into account in downstream analyses. Analyses that require genotype calls at the individual level, including haplotype-based analyses, should be performed with caution when using the data from the 1000 Genomes at *HLA* genes.

Our results have implications to studies that use SNP data from the 1000 Genomes in other genomic regions with high variability. Because *HLA* loci are the most polymorphic in the human genome, they represent a worst case scenario for mapping bias and subsequent allele frequency estimation errors. We found a significant correlation between SNP heterozygosity and the absolute difference in frequency between 1000 Genomes data and our gold standard. This suggests that in genome-wide studies, SNPs with high heterozygosities, and contained within regions with additional SNPs, have an increased chance of presenting poor frequency estimates.

## ACKNOWLEDGMENTS

This research was financially supported by São Paulo Research Foundation (FAPESP) scholarships #2012/22796-9 and #2013/12162-5 to DYCB and FAPESP research grant #12/18010-0 to DM. DM has an CNPq productivity grant 308167/2012-0.

